## Supplementary Figures for "Utilizing a dual endogenous reporter system to identify functional regulators of aberrant stem cell and differentiation activity in colorectal cancer"

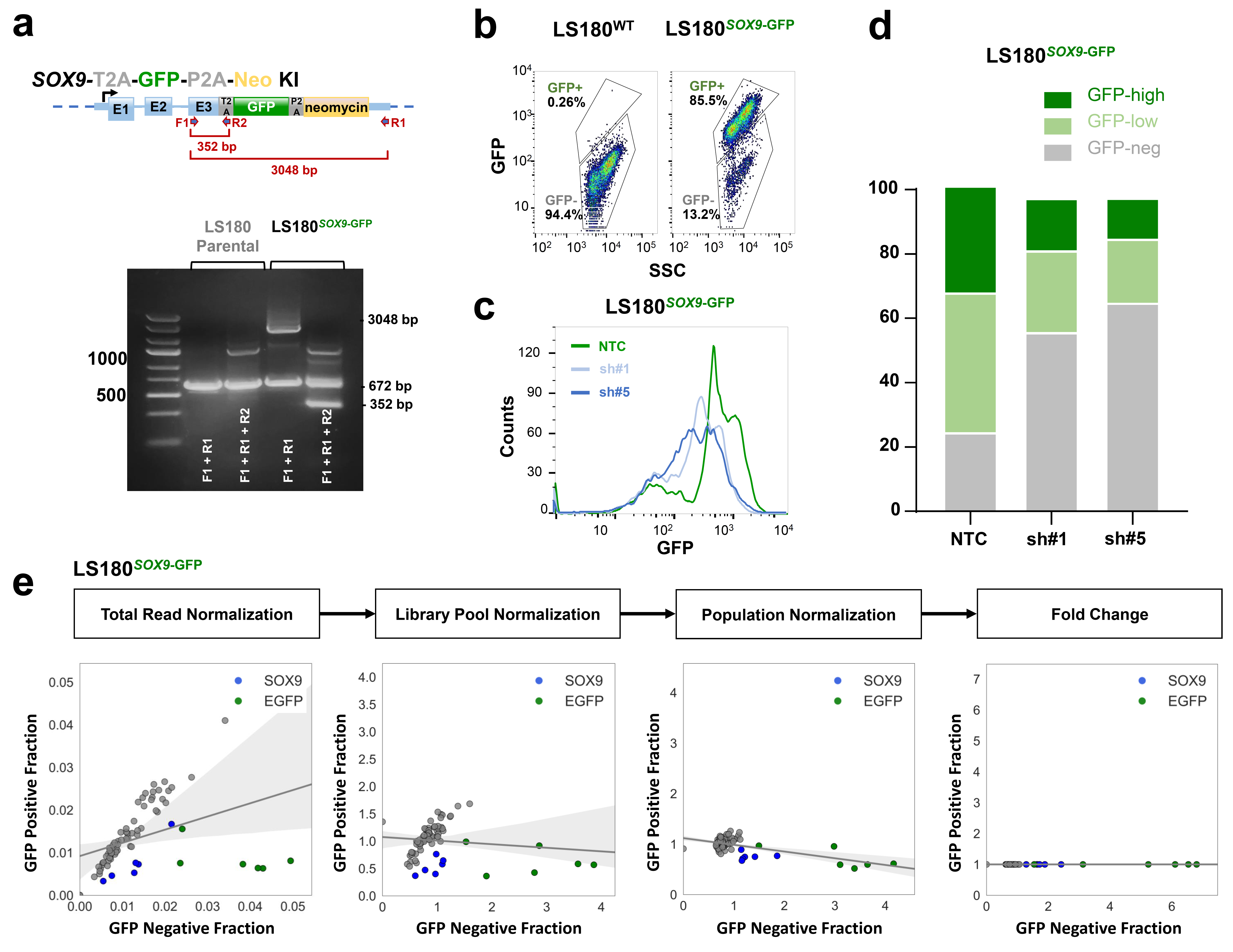
**Supplementary Figure 1.** Validation of the *SOX9* single reporter using custom CRISPR-Cas9 screen.

a. Validation primers F1, R1 and R2 are indicated by red arrows. Primers F1 and R1 amplifies a 672 bp genomic region in the unmodified parental cell line and a 3048 bp product after successful integration of reporter cassette. Primers F1 and R2 verify proper location and orientation of integrated cassette by amplifying a 352 bp product. PCR to validate cassette’s knock-in at the endogenous locus of SOX9. The expected size of correct knock-in is 3048 base-pairs.
b. Scatter plot of GFP/SSC showed gating scheme of LS180^SOX9-GFP^ single reporter versus LS180^WT^
c. Distribution of GFP intensity in LS180^SOX9-GFP^ single reporter upon SOX9 knock down with 2 different shRNAs compared to non-targeting shRNA (NTC).
d. Same as above
e. Scatter plot visualizing the effect of normalization on read count abundance in the top 2.5% (GFP Positive) and bottom 2.5% (GFP Negative) sorted fractions of the LS180SOX9-GFP single reporter cell line. Normalization of read counts by total read count (left panel), library pool (middle-left panel), and population normalization (middle-right panel), then calculation of fold change between the GFP Negative fraction compared to the GFP Positive fraction (right panel) are visualized.

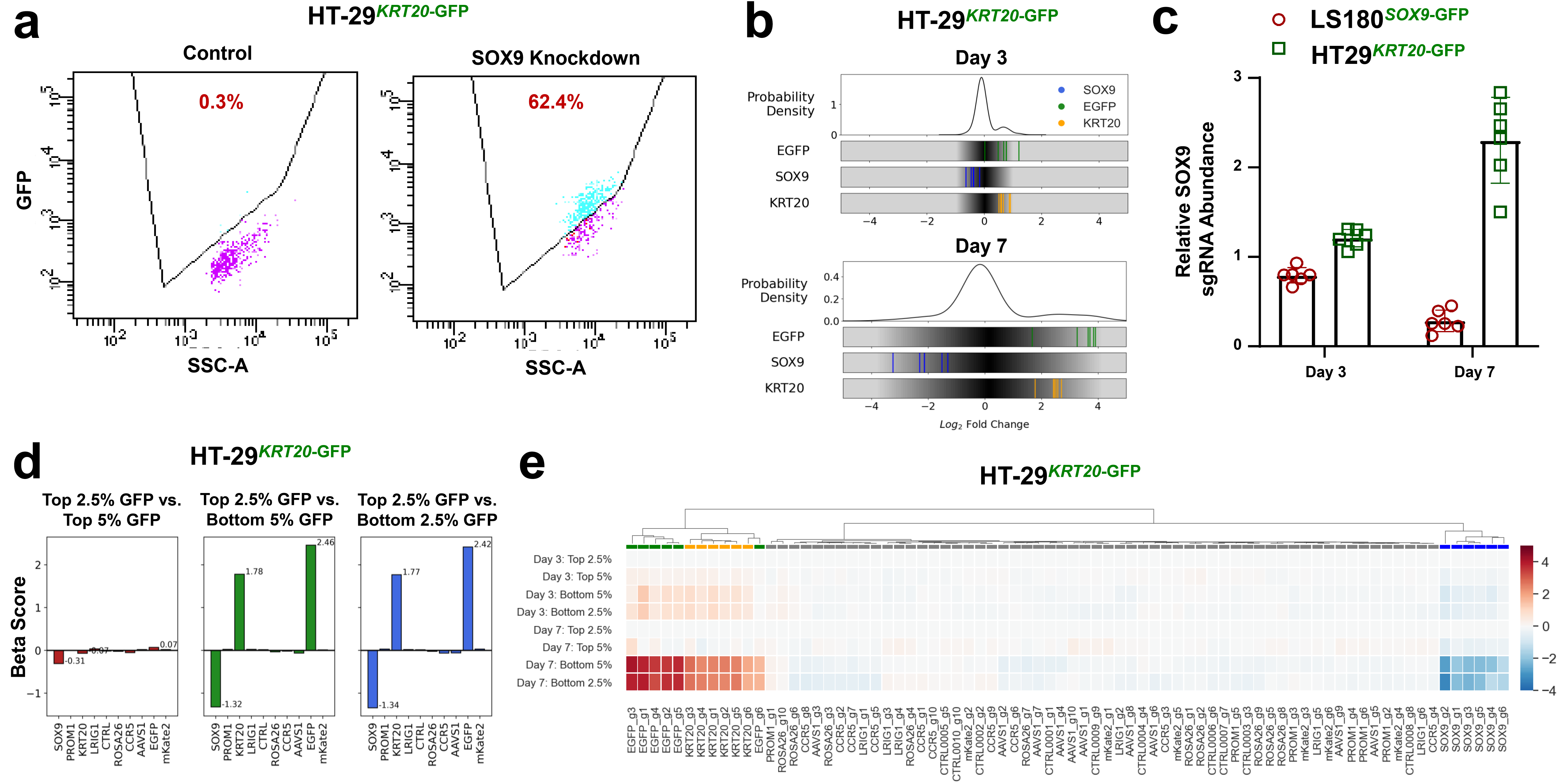
**Supplementary Figure 2.** Validation of *KRT20* single reporter system using custom CRISPR-Cas9 screen.

a. Flow analysis of HT29^KRT20-GFP^ single reporter upon SOX9 knockdown versus control.
b. Distribution of log2 fold change in the normalized abundance of all 76 sgRNAs in the focused CRISPR-Cas9 screen and GFP, SOX9, and KRT20 targeting sgRNAs comparing between the top 2.5% GFP positive and the bottom 2.5% GFP positive sorted cell fractions of the LS180-KRT20-EGFP cell line. The fold change of the normalized abundance of each sgRNA is shown at day 3 (top panel) and day 7 (bottom panel) post-library infection.
c. Relative abundance of 6 different sgRNA against SOX9 in LS180^SOX9-GFP^ single reporter and HT29^KRT20-GFP^ single reporter upon SOX9 knock-out.
d. Beta score of each gene in the epigenetic CRISPR-Cas9 screen (78 genes) comparing between the top 2.5% GFP fraction and the top 5% GFP fraction, bottom 5% GFP fraction, and bottom 2.5% GFP fraction of the HT29KRT20-EGFP cell line.
e. Hierarchical clustered heatmap of the fold change in the normalized abundance of the 76 sgRNAs in the focused CRISPR-Cas9 screen between each GFP-sorted fraction compared to either the top 2.5% GFP fraction at day 3 (top 4 rows) or day 7 (bottom 4 rows) post-library infection. KRT20 sgRNAs are represented by yellow markers. GFP sgRNAs are represented by green markers. SOX9 sgRNAs are represented by blue markers.

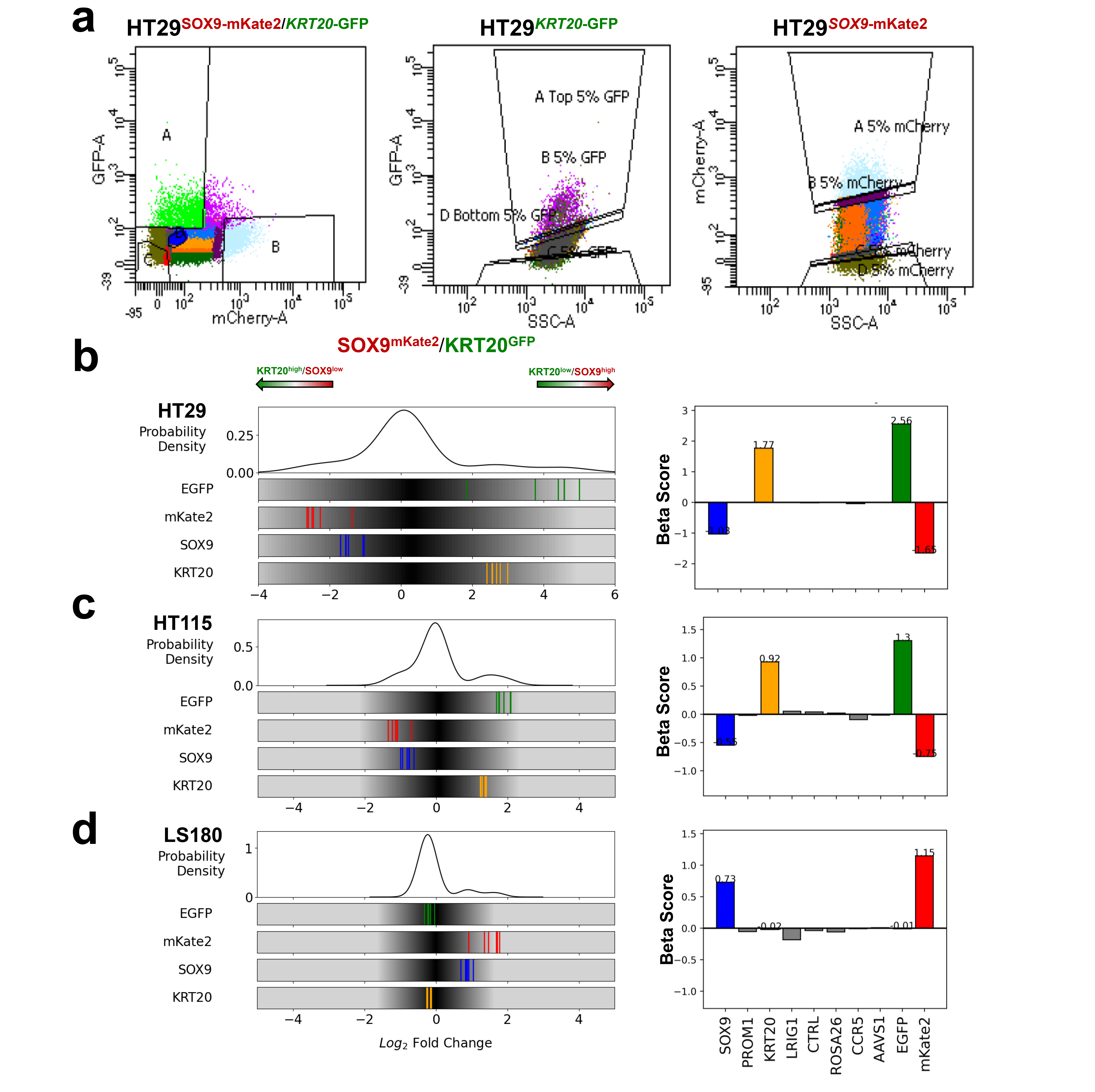

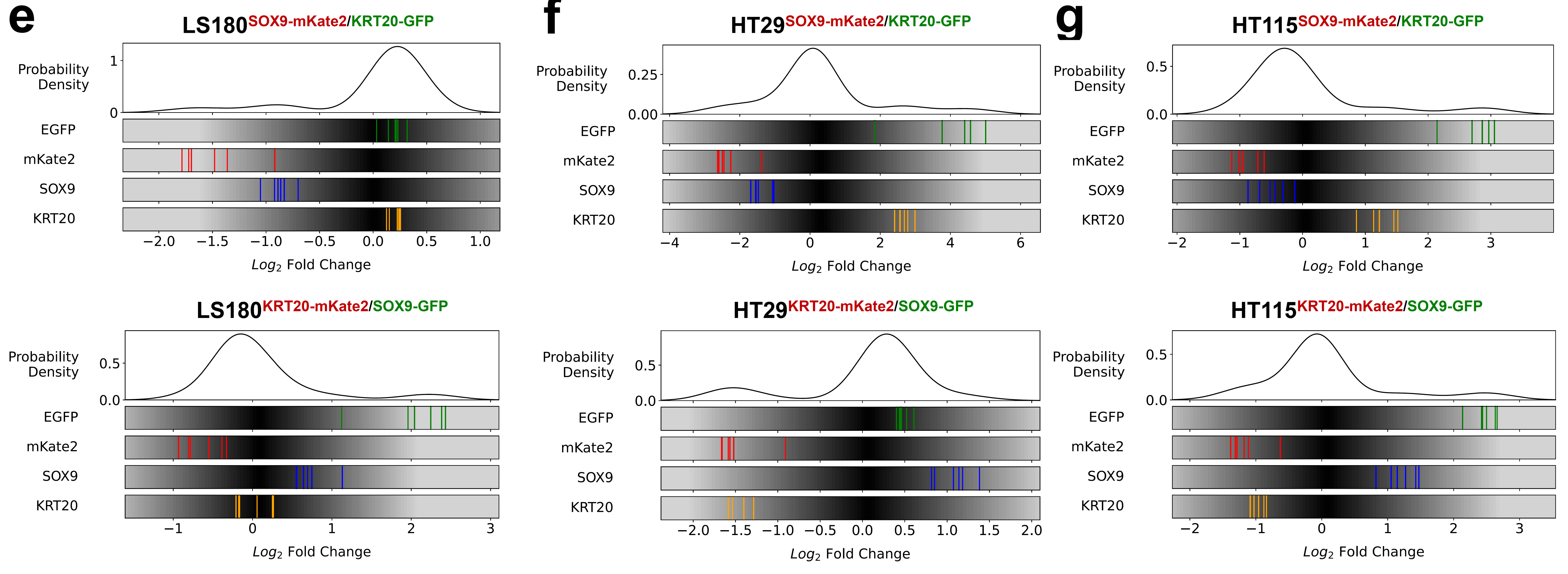

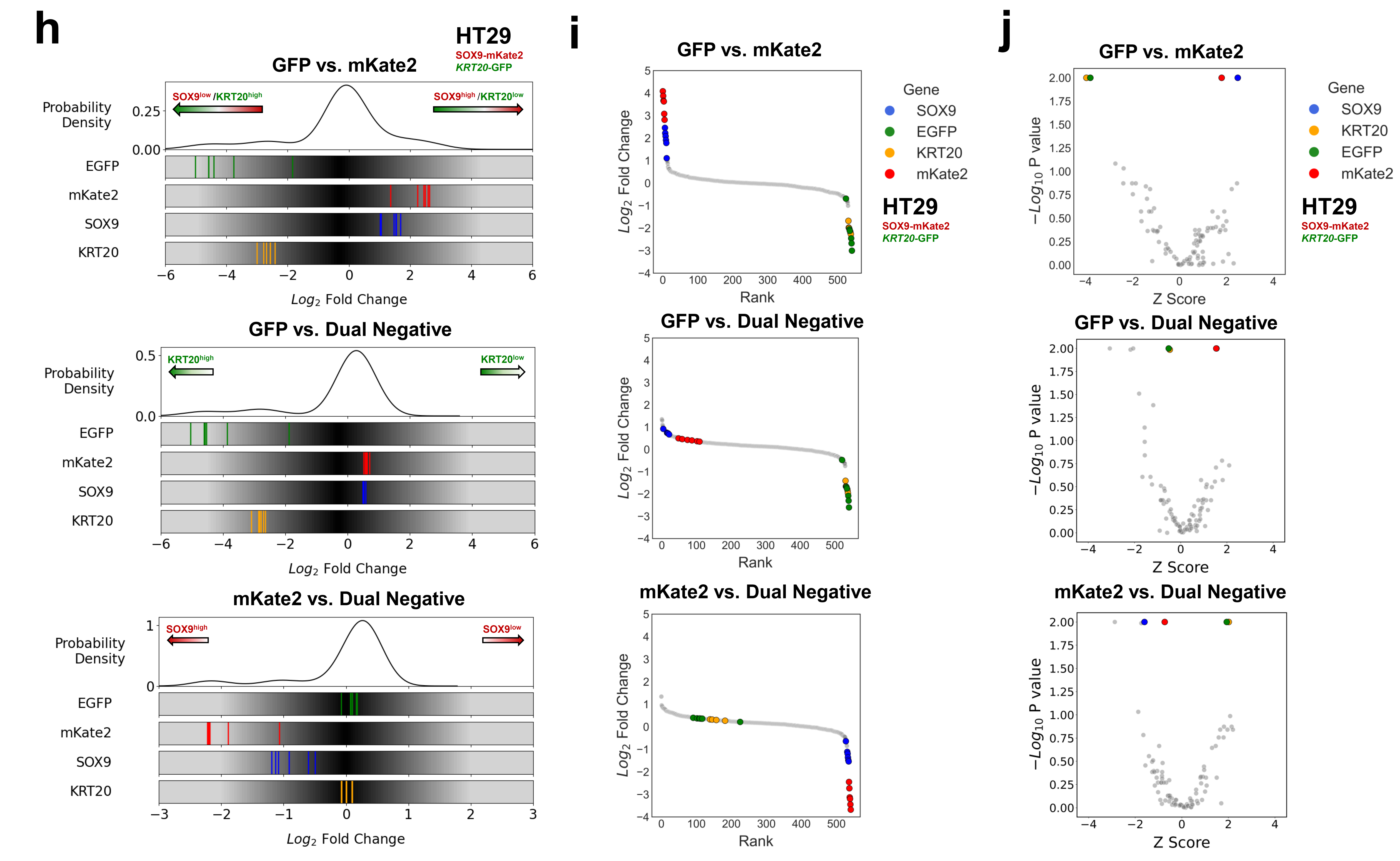
**Supplementary Figure 3.** Validation of dual endogenous reporter system using CRISPR-Cas9 screen.

a. After infection of focused CRISPR-Cas9 library (76 gRNAs) in the HT-29SOX9-mKate2/KRT20-GFP genome-edited cell line, cells were flow-sorted based on GFP and mCherry fluorescence intensity into 4 fractions: mKate2high/GFPhigh (D), mKate2high/GFPlow (B), mKate2low/GFPhigh (A), and mKate2low/GFPlow (C) (left panel). In comparison, the single endogenous differentiation GFP reporter system (middle panel) and the single endogenous stem cell mKate2 reporter system are flow-sorted into different fractions based on the fluorescence intensity of the single fluorophore.
b. Distribution of fold change of sgRNAs (left panel) and beta scores of gene essentiality (right panel) comparing between the mKate2low/GFPhigh fraction and the mKate2high/GFPlow fraction of the focused CRISPR-Cas9 screen in the HT29SOX9-mKate2/KRT20-GFP endogenous dual reporter cell line.
c. Distribution of fold change of sgRNAs (left panel) and beta scores of gene essentiality (right panel) comparing between the mKate2low/GFPhigh fraction and the mKate2high/GFPlow fraction of the focused CRISPR-Cas9 screen in the HT115SOX9-mKate2/KRT20-GFP endogenous dual reporter cell line
d. Distribution of fold change of sgRNAs (left panel) and beta scores of gene essentiality (right panel) comparing between the mKate2low/GFPhigh fraction and the mKate2high/GFPlow fraction of the focused CRISPR-Cas9 screen in the LS180SOX9-mKate2/KRT20-GFP endogenous dual reporter cell line

Distribution of log2 fold change of the sgRNA targeting the positive control genes and all sgRNAs in the focused CRISPR-Cas9 screen performed using the (e) LS180SOX9-GFP/KRT20-mKate2 and LS180SOX9-mKate2/KRT20-GFP genome-edited cell lines; (f) HT29SOX9-mKate2/KRT20-GFP, HT29KRT20-GFP/SOX9-GFP, (g) HT115SOX9-mKate2/KRT20-GFP and HT115KRT20-mKate2/SOX9-GFP genome-edited cell lines.

e. Distribution of fold change of sgRNAs comparing GFP versus mKate2 (top), GFP versus dual negative (middle), and mKate2 versus dual negative (bottom) in the HT29SOX9-mKate2/KRT20-GFP endogenous dual reporter cell line
f. Distribution of fold change of sgRNAs versus Rank when comparing GFP versus mKate2 (top), GFP versus dual negative (middle), and mKate2 versus dual negative (bottom) in the HT29SOX9-mKate2/KRT20-GFP endogenous dual reporter cell line
g. Distribution of fold change of sgRNAs versus Z-score when comparing GFP versus mKate2 (top), GFP versus dual negative (middle), and mKate2 versus dual negative (bottom) in the HT29SOX9-mKate2/KRT20-GFP endogenous dual reporter cell line
h. Distribution of sgRNA log2 fold change of the focused library (76 gRNAs) in the dual reporter system. Log2 fold change of normalized sgRNA read counts is shown by comparing the GFP positive and mCherry positive sorted cell fractions (top), GFP positive and Dual Negative fraction (middle), and mCherry vs. Dual Negative fraction (bottom) in the HT-29SOX9-mKate2/*KRT20*-GFP cell line.
i. Ranked log2 fold change plot of the epigenetic CRISPR-Cas9 screen (542 gRNAs) using the dual reporter system in the HT-29SOX9-mKate2/*KRT20*-GFP cell line.
j. Volcano plot of the epigenetic CRISPR-Cas9 screen (88 genes) using the dual reporter system in the HT-29SOX9-mKate2/*KRT20*-GFP cell line. The *x* axis shows the *Z* score of gene-level FC (mean log2(fold change) for all sgRNAs targeting the same gene). The y axis shows the gene-level p values generated by MaGeCK Maximum Likelihood Estimation.

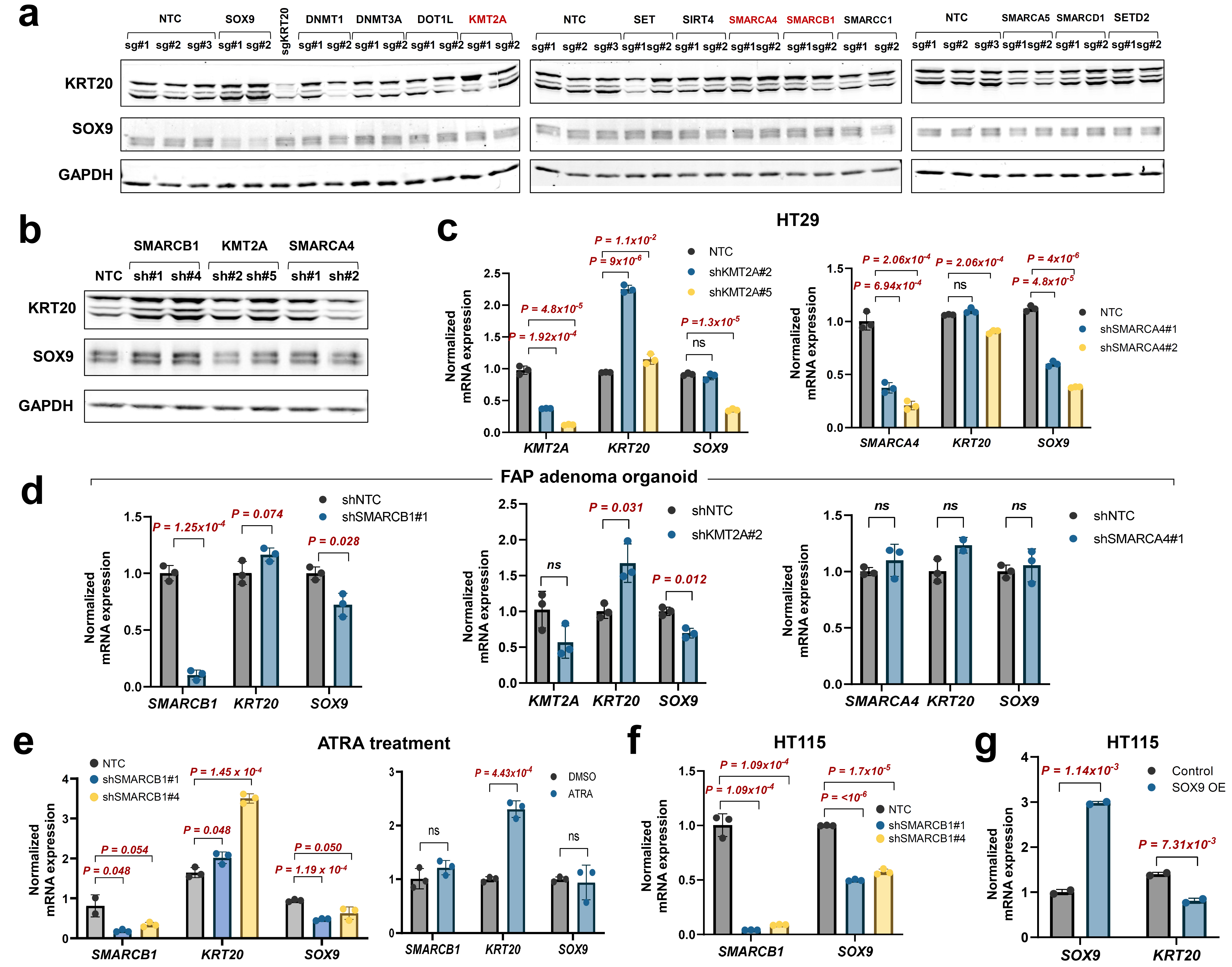
**Supplementary Figure 4.** Validation of dual reporter hits using CRISPR knockout and shRNA-mediated knockdown in different CRC models.

a. Immunoblot of KRT20, SOX9, and GAPDH (loading control) in HT-29 CRC cells following CRISPR-Cas9 knockout of 12 top-scoring genes from dual reporter screen.
b. Immunoblot of KRT20, SOX9, and GAPDH in HT-29 CRC cells engineered to knockdown (KD) SMARCB1, KMT2A, and SMARCA4 with 2 different hairpins.
c. Normalized mRNA expression levels of *KMT2A*, *SMARCA4*, *KRT20*, and *SOX9* in HT-29 CRC cells with KMT2A and SMARCA4 KD by RT-PCR
d. Normalized mRNA expression levels of *SMARCB1*, *KMT2A*, *SMARCA4*, *KRT20*, and *SOX9* in adenoma organoids from a patient with familial adenomatous polyposis (FAP) engineered to knockdown SMARCB1, KMT2A, and SMARCA4.
e. Normalized mRNA expression levels of *SMARCB1*, *KRT20*, and *SOX9* in SMARCB1 KD (left) and parental (right) HT-29 cells with ATRA treatment at 10 uM at 48 hr.
f. Normalized mRNA expression levels of *SMARCB1* and *SOX9* in SMARCB1 KD HT-115 CRC cells by RT-PCR
g. Normalized mRNA expression levels of SOX9 and KRT20 in HT-115 CRC cells engineered for inducible expression of SOX9 cDNA.

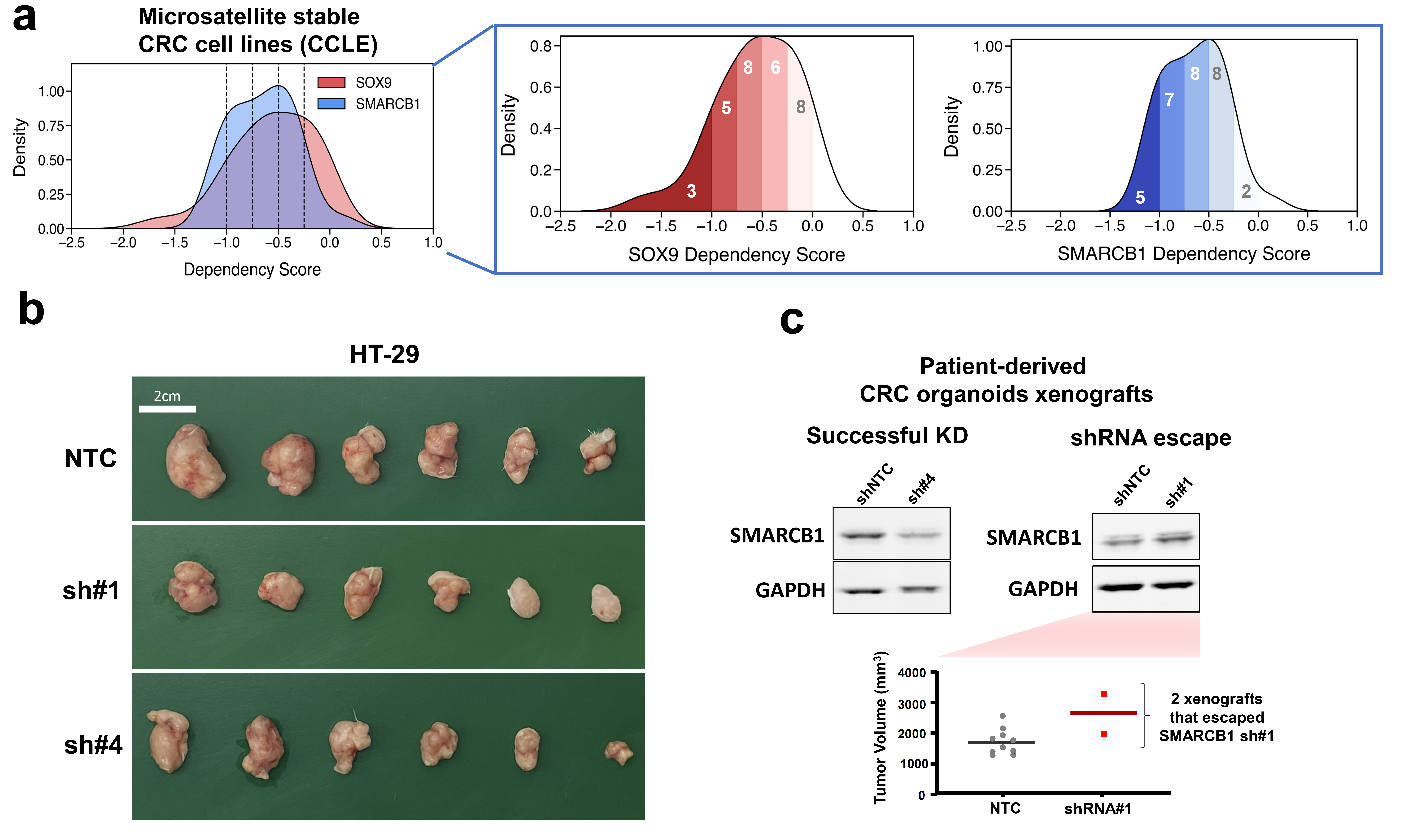

**Supplementary Figure 5.** SMARCB1 is a dependency in CRC.

a. Distribution of dependency scores across microsatellite stable CRC cell lines (n=31) for the positive control gene of stemness used in the endogenous reporter system (SOX9) and the most promising candidate dependency in CRC for validation (SMARCB1), annotated using the number of cell lines where the target gene is below each dependency score threshold.
b. Photograph of HT29 xenograft tumors at experimental endpoint.
c. Immunoblot showing successful SMARCB1 KD (left) and shRNA escape in patient-derived CRC organoid xenografts; tumor volumes for two xenografts that escaped shRNA KD and nontargeting control (NTC) xenografts at Day 32.

**Supplementary Tables**

**Supplementary Table 1.** Sequences of sgRNA and homology arm for SOX9 and KRT20 knock-in

**Supplementary Table 2.** List of 78 epigenetic regulators from 5 families and their associated drugs

**Supplementary Table 3.** List of 76 sgRNAs and 154 shRNAs for control libraries and 542 sgRNAs for the epigenetic library

**Supplementary Table 4.** List of sequences used in the validation

**Supplementary Table 5**. List of antibodies used for the western blot
